# *Fbxo2*^*VHC*^ mouse and embryonic stem cell reporter lines delineate *in vitro*-generated inner ear sensory epithelia cells and enable otic lineage selection and Cre-recombination

**DOI:** 10.1101/352716

**Authors:** Byron H. Hartman, Robert Böscke, Daniel C. Ellwanger, Sawa Keymeulen, Mirko Scheibinger, Stefan Heller

## Abstract

While the mouse has been a productive model for inner ear studies, the lack of highly specific genes and tools have presented challenges, specifically for *in vitro* studies of otic development, where innate cellular heterogeneity and disorganization increase the reliance on lineage-specific markers. To address this challenge in mice and embryonic stem (ES) cells, we targeted the lineage-specific otic gene *Fbxo2* with a multicistronic reporter cassette (Venus/Hygro/CreER = VHC). In otic organoids derived from ES cells, *Fbxo2^VHC^* specifically delineates otic progenitors and inner ear sensory epithelia. In mice, Venus expression and CreER activity reveal a cochlear developmental gradient, label the prosensory lineage, show enrichment in a subset of type I vestibular hair cells, and expose strong expression in adult cerebellar granule cells. We provide a toolbox of multiple spectrally distinct reporter combinations to the community for studies that require use of fluorescent reporters, hygromycin selection, and conditional Cre-mediated recombination.

**SUMMARY STATEMENT:** A multifunctional *Fbxo2*-targeted reporter in mice and stem cells was developed and characterized as a resource for inner ear studies, along with a toolbox of plasmids to facilitate the use of this technique for other users.

## INTRODUCTION

The sensory organs of the inner ear contain highly specialized mechanosensory epithelia, essential for hearing and balance. The five vestibular organs and the auditory organ, the organ of Corti, originate from the otic vesicle during embryonic development and are comprised of various types of sensory hair cells and supporting cells. Auditory and vestibular hair cells are innervated respectively by spiral ganglion or vestibular ganglion neurons, which also originate in the otic vesicle. The inner ear offers an excellent developmental system to study mechanisms of lineage specification, cell fate determination, patterning, and organogenesis (Wu and Kelley, 2012). The mouse has been a highly productive model system for inner ear studies due to advances in mouse genetics and the relative similarity of the mouse inner ear to that of humans and other mammals. Studying the murine otic lineage, however, has been challenging because very few genes are expressed exclusively in the inner ear. Most genes that have been found to be important for inner ear development play important roles in development of other organs as well (Dahl et al., 1997; Tadjuidje and Hegde, 2013; Torban and Goodyer, 2009). Also, some genes that have been used as favored markers or exploited as transgenic tools for the developing inner ear are expressed transiently in the otocyst, which limits their use (Chatterjee et al., 2010; Wu and Kelley, 2012).

Considerable progress has been made in ES cell models of otic development, where stem cells can be coaxed into the otic sensory lineage through manipulation of signaling to eventually produce otic sensory cell types (DeJonge et al., 2016; Ealy et al., 2016; Koehler et al., 2013; Koehler et al., 2017; Li et al., 2003; Oshima et al., 2010). However, stem cell differentiation is inherently heterogeneous and, without highly specific and persistent otic lineage markers, it is currently necessary to assay for multiple markers at different time points in order to infer the identity of potential otic cells (Ealy et al., 2016; Koehler et al., 2013; Oshima et al., 2010; Schaefer et al., 2018). This limitation is particularly the case for identification of otic progenitors and supporting cells, which lack the more distinguishing cytological features and expression profiles of hair cells. A second major challenge for the field is that there are limited mouse lines available for driving expression of Cre recombinase specifically in the inner ear (Cox et al., 2012).

We recently isolated and characterized mouse otic sensory lineage-specific genes and identified *Fbxo2* as the most differentially expressed otic gene at embryonic day 10.5 (E10.5) (Hartman et al., 2015). *Fbxo2* encodes the ubiquitin ligase subunit F-box 2 (Fbx2), which we found to be strongly expressed in the developing inner ear starting at E10.5. Consistent with our previous study, *Fbxo2* mRNA is
specifically detectable in the mouse inner ear at E14.5, with little or no signal in other tissues of the embryo (Fig. 1A, Eurexpress database assay 004104, Diez-Roux et al. (2011)). At later stages of embryonic development Fbx2 remains highly specific to the inner ear and is enriched in the auditory and vestibular sensory epithelia cells including sensory hair cells and supporting cells (Fig. 1B-B`, and Hartman et al. (2015)). These data suggested that at both the transcript and protein level *Fbxo2* stands out as a highly specific and robust marker of the otic sensory lineage, which motivated us to focus on this gene.

**Figure 1.**
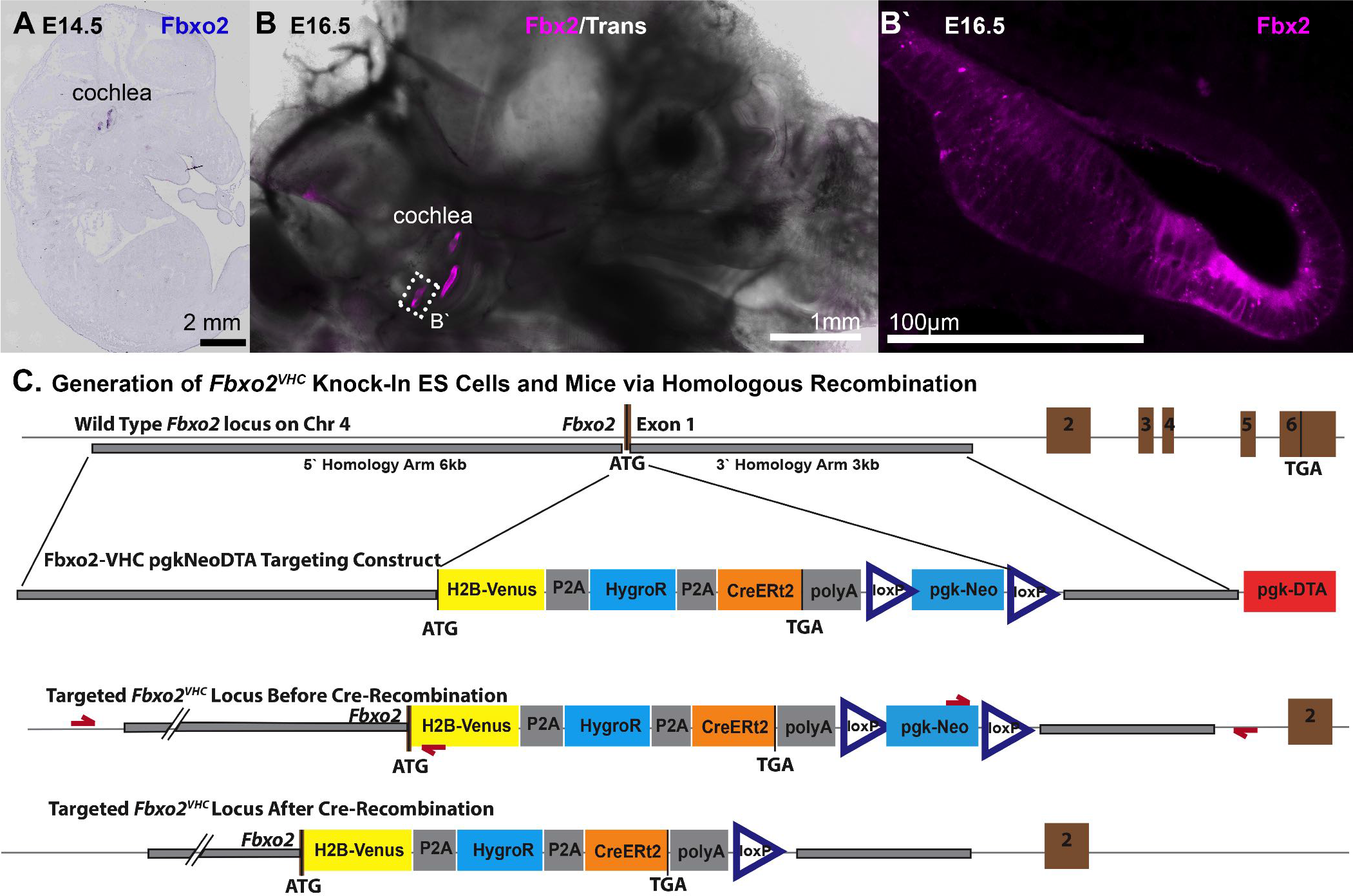
Generation of *Fbxo2*^*VHC*^ multicistronic reporter and mouse knock-in allele. A) *Fbxo2 mRNA in situ* hybridization in E14.5 mouse (Eurexpress database) shows strong enrichment specifically in the developing inner ear. B) Immunolabeling for Fbx2 protein in a thick vibratome section of E16.5 mouse head shows high expression specifically in the developing inner ear. B`) Higher magnification view of Fbx2 immunolabeling in the basal turn of the cochlea shows expression in the cytoplasm of sensory hair cells and supporting cells. C) Schematic showing the generation of *Fbxo2*^*VHC*^ mice and ES cells. The wild type *Fbxo2* locus on Chr 4 contains six exons, with the ATG start site for Fbx2 in Exon 1. The Fbxo2VHCpgkNeoDTA targeting vector was constructed with homology arms from the indicated regions of Chr 4, flanking a “knock-in” cassette consisting of the VHC reporter followed by floxed pgk-Neo for positive selection. A pgk-DTA cassette was positioned downstream of the 3` homology arm, for negative selection of off-target genomic insertions. The third line shows a representation of the correctly targeted *Fbxo2*^*VHC*^ locus, essentially replacing Exon 1 of *Fbxo2* with the VHC cassette, and still carrying the pgk-Neo, which was essential for selection of ES cell colonies. Redarrows represent the location of primers for PCR screening of ES cell clones. Verified clones were used to make chimeric mice and two clones were transmitted to offspring to produce viable mouse lines. The bottom schematic is the *Fbxo2*^*VHC*^ locus after removal of the pgk-Neo cassette in the germ line of mice through further out-crossing to Eiia-Cre transgenic mice. The final *Fbxo2*^*VHC*^ locus, with pgk-Neo removed, was then isolated from the Eii-Cre transgene and maintained in heterozygous offspring over at least five generations of back-crossing onto the FVB-NJ background.

Fbx2 (also called Fbs1, NFB42, and OCP1) is a sugar chain specific F-box protein that was previously identified in the adult guinea pig organ of Corti where it was hypothesized to form a complex with Skp1 and interact with connexin 26 (Cx26), an essential cochlear gap junction protein (Henzl et al., 2001; Henzl et al., 2004; Thalmann et al., 2003; Thalmann et al., 1997). Fbx2 binds to high-mannose, asparagine-linked glycoproteins that are associated with the endoplasmic reticulum, and is believed to facilitate their degradation through the ubiquitin proteasome pathway (Yoshida et al., 2002; Yoshida et al., 2003). Fbxo2^−/−^ mice have normal inner ear morphology and hearing prior to two months of age; thereafter, they exhibit age-related cochlear degeneration and hearing loss (Nelson et al., 2007). This suggests that in the adult inner ear Fbx2 plays an essential role, presumably in protein quality control. It is interesting that *Fbxo2* is expressed robustly from early otic development, but the knockout does not display a detectable phenotype until months later. *Fbxo2* is also expressed in the adult brain where there is evidence it may influence NMDA subunit distribution as well as attenuate Alzheimer’s disease amyloidosis by mediating ubiquitination of beta-secretase (Atkin et al., 2014; Atkin et al., 2015; Erhardt et al., 1998; Gong et al., 2010). It is currently not known when *Fbxo2* becomes expressed in the brain or which populations of neurons most highly express this gene. The lack of substantial expression signal for *Fbxo2* in other embryonic tissues indicates that levels in other tissues are much lower than in the inner ear, or that expression is initiated at later embryonic or postnatal stages in other tissues.

We sought to exploit the otic-specific expression of *Fbxo2* by generating embryonic stem cells and a mouse line in which *Fbxo2*-expressing cells can be identified, isolated, and genetically manipulated for studies of inner ear development and function. Such a reporter system could serve as a tool to solve the issue of cell heterogeneity in stem cell-based model systems of inner ear development, including somatic direct programming and directed stem cell differentiation approaches, which are currently reliant on relatively non-specific markers and tools for identifying and accessing newly generated inner ear cell types. To achieve these goals, we generated a tricistronic reporter cassette (VHC) consisting of the bright fluorescent nuclear protein H2B-Venus (V), a hygromycin resistance gene (HygroR = H), and the CreERt2 gene (C) that was knocked into the *Fbxo2* locus. *Fbxo2*^*VHC/WT*^ ES cells displayed specific and early otic progenitor cell expression of the Venus reporter that persisted when the otic progenitors differentiated into supporting cells and hair cell-like cells. In *Fbxo2*^*VHC/WT*^ mice, H2B-Venus expression is moderate in the early otic vesicle and becomes strong and concentrated throughout the floor of the developing cochlear duct and later in supporting cells and hair cells of the organ of Corti. In neonatal and adult mice, H2B-Venus expression is also detectable in subsets of other cochlear cell types. Outside the inner ear, we identified strong reporter gene expression in adult cerebellar granule cells. We further validated the reporter system’s selection marker for hygromycin as well as the use of tamoxifen-induced and CreERt2-mediated recombination. In summary, the *Fbxo2*^*VHC*^ reporter system presented here exploits the embryonic specificity of *Fbxo2* expression to enable specific delineation and manipulation of otic (pro-) sensory cells in culture systems and mice.

## RESULTS

### Generation of multicistronic reporter cassettes and *Fbxo2*^*VHC*^ knock-in ES cells

We developed a multifunctional reporter system for 1) expression of a bright fluorescent protein for imaging, 2) conferring drug-resistance for selection, and 3) enabling inducible Cre-mediated recombination for lineage tracing and conditional mutagenesis when combined with LoxP mouse models. Viral 2A peptides are small “self-cleaving” amino acid chains that enable expression of multiple proteins from a single transcript by mediating a ribosomal skipping of the last peptide bond at the C-terminus (Szymczak and Vignali, 2005). We designed and constructed a tricistronic reporter cassette using the P2A peptide, from porcine teschovirus-1, which has been shown to have high cleavage efficiency in human cell lines, mice, and zebrafish (Kim et al., 2011). The resulting H2B Venus-P2A-HygroR-P2A-CreER cassette (abbreviated VHC) encodes expression of three products from a single transcript (Fig. S1A). The first product, H2B-Venus, is a carboxyl-terminal fusion of human histone H2B with the yellow fluorescent Venus protein. Nuclear located H2B fluorescent fusion proteins have been widely used for studies of cell cycle, apoptosis, and chromatin dynamics (Fraser et al., 2005; Kang et al., 2013; Nowotschin et al., 2009). The second cDNA, HygroR encodes hygromycin phosphotransferase that confers resistance to hygromycin B for selection of cells in culture (Blochlinger and Diggelmann, 1984; Giordano and McAllister, 1990). HygroR has been used effectively in 2A mediated bicistronic expression for selection of ES cells following dual recombinase mediated cassette exchange (Osterwalder et al., 2010). The third cDNA in the VHC cassette is CreERt2, which enables inducible Cre-LoxP recombination for lineage tracing (Feil et al., 1997; Matsuda and Cepko, 2007). The sequence of the three proteins in the VHC cassette was designed to minimize possible issues associated with the 2A “tag” that remains on the carboxyl terminus of any protein upstream of the 2A peptide (Szymczak and Vignali, 2005). We therefore placed CreERt2 in the last position to ensure that the estrogen receptor fusion remains unmodified. A similar strategy has been used successfully in knock-in mice for multicistronic 2A-peptide cassettes expressing CreERt2 downstream of other proteins (Imuta et al., 2013). The VHC cassette was designed with two P2A peptides of identical amino acid sequence with slightly different cDNA sequences to reduce repetitive sequences (Fig. S1A). Upstream of the VHC translation initiation codon, we included a Kozak consensus sequence (GCCACCATG (Kozak, 1987)) overlaid with an in-frame eight-nucleotide rare restriction endonuclease recognition site (Fse-1, GGCCGGCC (Nelson et al., 1990)) to enable scarless assembly of promoter sequences or homology arms to the 5`-end of the reporter cassette (Fig. S1A).

We tested the VHC cassette in cell culture to verify functional performance of the three reporter proteins (Fig. S1 B-G). The VHC cassette was cloned downstream of the ubiquitous cytomegalovirus immediate early enhancer-chicken beta-actin hybrid (CAG) promoter (Fig. S1B) (Okabe et al., 1997; Sakai et al., 1995). Embryonic fibroblasts generated from Ai14 Cre reporter mice (Madisen et al., 2010) were transfected with pCAG-VHC, treated with 4-hydroxytamoxifen (4OHT) and HygroB, and then assayed for expression of Venus. Yellow fluorescence from Venus expression was bright and localized to the nucleus and Venus-positive cells treated with 4OHT expressed the tdTomato Cre reporter (Fig S1D-F). HygroB treatment resulted in a substantial increase in the fraction of Venus expressing cells (Fig. S1G). To generate the targeting construct to knock-in the VHC cassette to the *Fbxo2* locus, we cloned the VHC cassette into the PGKneolox2DTA.2 plasmid (Fig. S1H) (Hoch and Soriano, 2006) that contains a neomycin selection cassette flanked by LoxP sites (to allow subsequent removal by Cre recombination) and a PGKdtabpA (for negative selection of non-homologous insertion). We then generated knock-in *Fbxo2*^*VHC*^ ES cells by cloning homology arms flanking the start site of the *Fbxo2* gene into the VHC targeting vector and performed homologous recombination and selection of neomycin-resistant ES cell colonies (Fig. 1C).

We built and validated three additional multicistronic cassettes as part of a toolbox of plasmids to facilitate use of this approach in a variety of systems and (Fig. S2). We generated four versions of the cassette with alternative fluorescent proteins, specifically H2B-Venus (VHC), H2B-Emerald (EHC), H2B-tdTomato (tdTHC), and H2B-Teal/mTFP1 (mTHC) (Fig. S2A). These four fluorescent proteins are spectrally distinct and among the brightest in their respective classes (Day and Davidson, 2009). The properties of these fluorescent proteins are well characterized and they can be excited by common laser lines and distinguished by their excitation and emission spectra (Fig S2B-C). Three plasmids were generated for each of the multicistronic cassettes: a promoter-less expression plasmid, a CAG promoter plasmid for constitutive expression, and a PGKneolox2DTA.2 plasmid for generation of targeting vectors by addition of homology arms (Fig. S2H).

### Venus marks *Fbxo2*^*VHC*^ ES cell-derived inner ear sensory epithelia cells

Here we show proof-of-principle that the *Fbxo2*^*VHC*^ reporter is useful in specifically delineating otic cells produced from ES cells by directed differentiation. *Fbxo2*^*VHC/WT*^ ES cell aggregates were sequentially treated with small molecules and recombinant proteins to manipulate Bmp, TGFβ, Fgf, and Wnt signaling pathways, according to a previously established three-dimensional culture system for otic sensory differentiation (DeJonge et al., 2016; Koehler and Hashino, 2014). This protocol promotes stepwise induction of non-neural and preplacodal ectoderm, followed by development of otic vesicle-like structures by day 12-14, which ultimately contain mechanosensory hair cells and supporting cells by day 16 or later (Koehler and Hashino, 2014). In *Fbxo2*^*VHC/WT*^ ES cell-derived aggregates at day 12 of the otic differentiation protocol, epithelial cysts co-express Venus and Pax2 (Fig. 2A-A`). Pax2 is expressed in the early otic placode and synergizes with Pax8 for specification of inner ear (Bouchard et al., 2010; Burton et al., 2004; Hans et al., 2004; Nornes et al., 1990). During early otic morphogenesis at E10.5 in the mouse embryo Pax2 and Fbx2 are coexpressed in the otic vesicle (Hartman et al., 2015). We observed *Fbxo2*^*VHC/WT*^ aggregates at 17 days *in vitro* containing numerous epithelial cysts (organoids) of Venus-expressing cells (Fig. 2B). Fbx2 immunoreactivity is specifically found in organoids of Venus-expressing epithelial cells within *Fbxo2*^*VHC/WT*^ ES cell aggregates, confirming specificity of the knock-in reporter (Fig. 2B-B`, Movie 1). Early hair cell markers MyosinVI (Myo6), MyosinVIIa (Myo7a) and Gfi1 are expressed in the lumenal Venus-positive cells while the cells in the basal layer express the supporting cell marker Jagged1 (Jag1) (Fig. 2C-E``). In day-26 organoids, Calretinin-positive hair cell-like cells are associated with nerve endings projecting from Tuj1-labeled neurons (Fig. 3A-B``, Movie 2). Sox2 expression is maintained in hair cell-like cells and supporting cells, consistent with previous reports (Koehler et al., 2013) (Fig. 3C-C`). The hair cell-like cells feature stereocilia on the lumenal surface that were enriched with the hair bundle protein Espin (Fig. 3D-F, Movie 2). Overall, in organoids at days 17 and 26, H2B-Venus is expressed in hair cell-like cells and supporting cells and adjacent epithelial cells. Its expression comprehensively delineates otic cell populations. Within putative hair cell and supporting cell populations the reporter shows intensity variations in domains and patches, somewhat reminiscent of pattern formations of inner ear organogenesis (Fig. 3, Movie 3 (Brigande et al., 2000; Groves and Fekete, 2012)).

**Figure 2.**
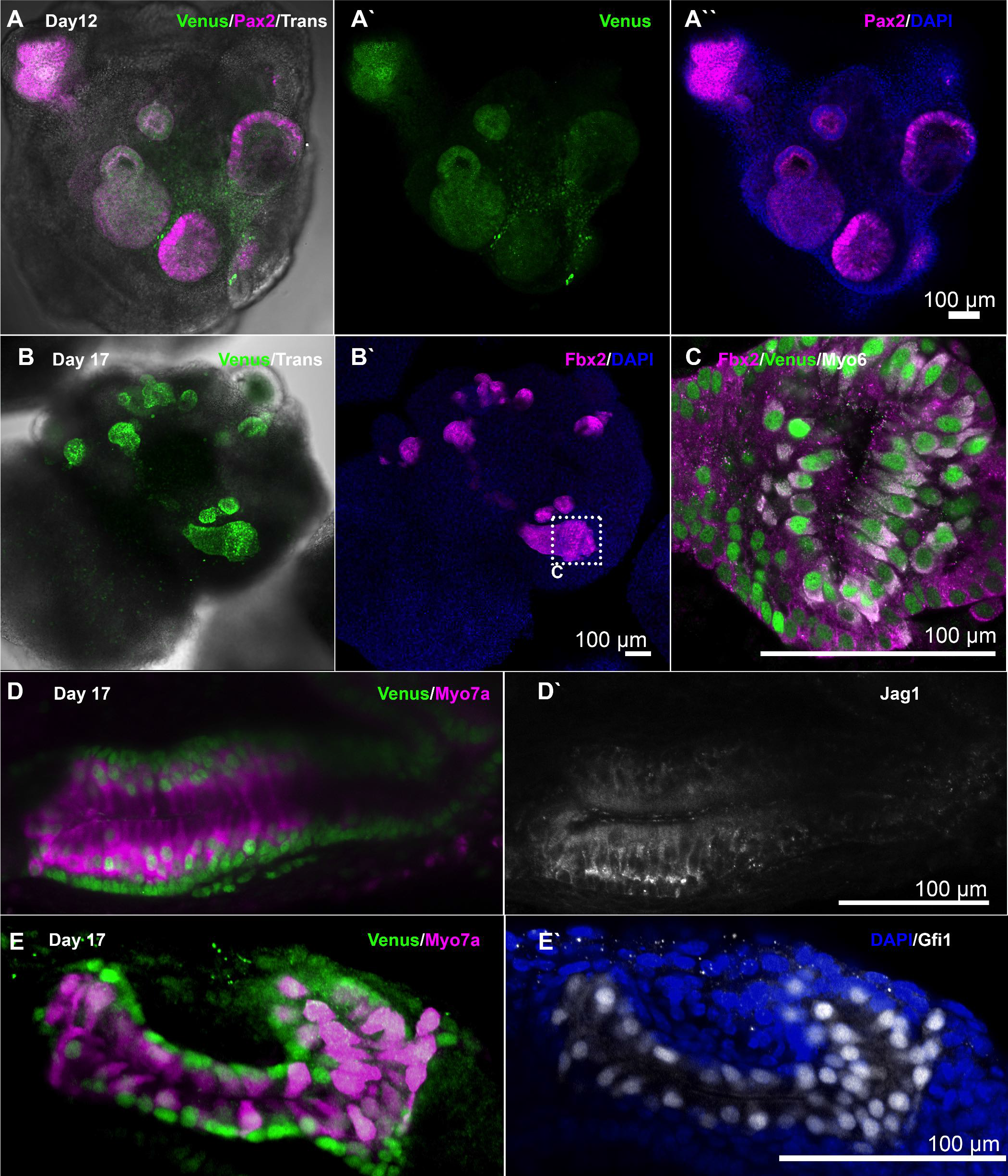
Venus expression in developing sensory epithelia in otic organoids produced from *Fbxo2*^*VHC*^ ES cells. *Fbxo2*^*VHC*^ ES cell aggregates at day 12 (A-A``) or day 17 (B-E`) of the *in vitro* inner ear differentiation protocol. Aggregates were fixed and prepared for whole-mount immunolabeling with antibodies to the indicated markers. A-A``) Projection views of a confocal micrograph z-stack taken of aday 12 aggregate containing organoids composed of epithelial cells that co-express Venus and the early otic vesicle marker Pax2. B-B`) Example of an aggregate at day 17 containing otic organoids marked by co-expression of H2B-Venus and Fbx2. C). Higher magnification view of a single optical section taken from the boxed region marked in panel B, showing co-expression of Fbx2 and Venus in epithelial cells. A small luminal space can be seen in the middle, lined with cells that express the hair cell marker Myo6. D-D`) An example of Venus expression in a day 17 organoid with Venus and Myo7a-positive cells with hair cell morphology, surrounded by Venus and Jag1-expressing epithelial cells resembling supporting cells. E-E`) A day 17 inner ear organoid composed of Venus-expressing cells including Myo7a and Gfi1 co-expressing hair cells.

**Figure 3.**
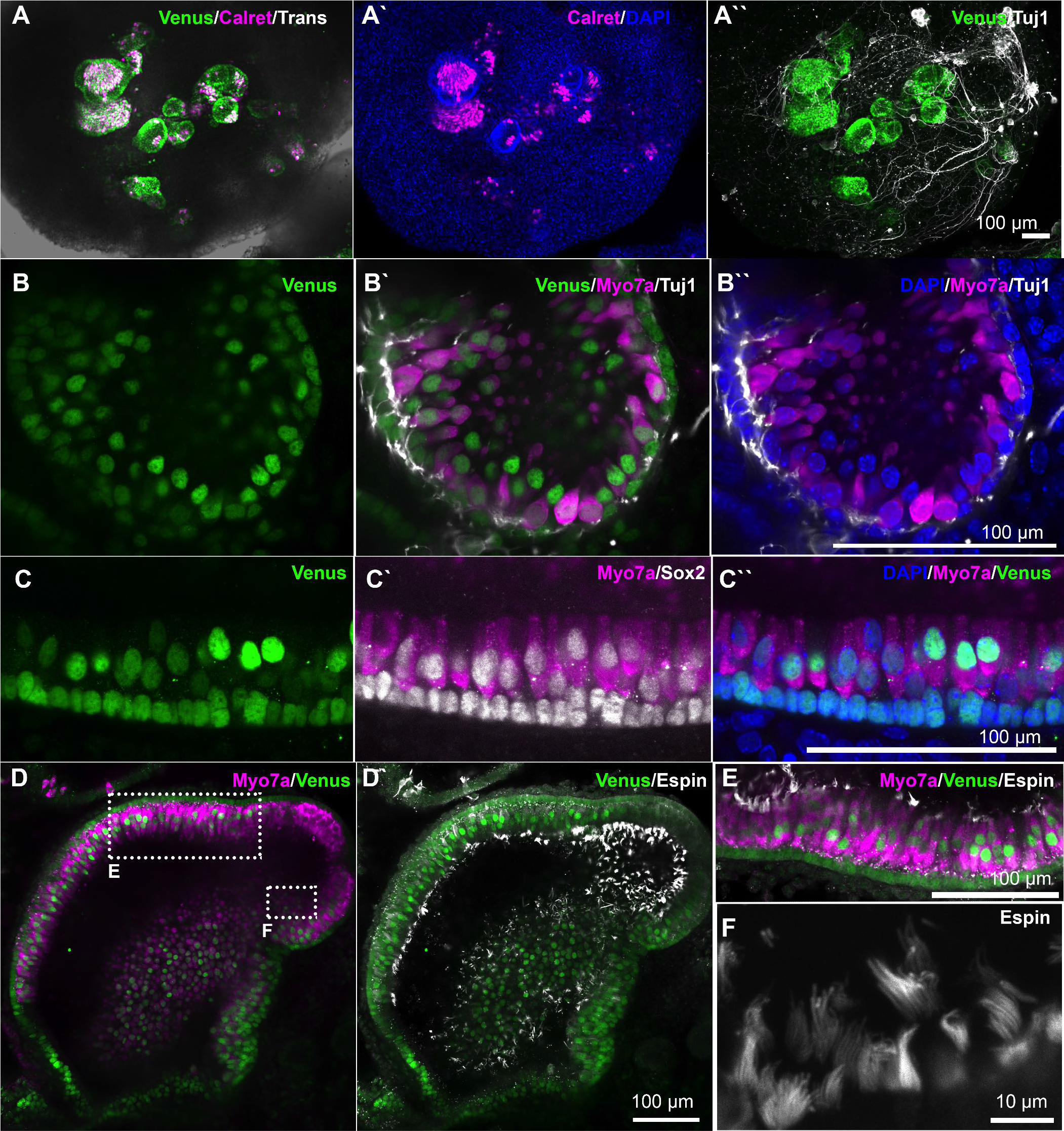
Venus expression in hair cells and supporting cells of day 26 *Fbxo2*^*VHC*^ ES cell otic organoids. *Fbxo2*^*VHC/WT*^ ES cell aggregates after 26 days of *in vitro* inner ear differentiation protocol. Aggregates were fixed and prepared for whole-mount immunolabeling with antibodies to the indicated markers. A-A``) Epithelial cysts demarcated by Venus expression contain Calretinin-positive hair cells while Tuj1 immunoreactivity labels neurons with ganglion-like morphology that feature projections associated with the cysts. B-B`) A projected view of the inside of a cyst that shows Tuj1-labeled nerve endings associated with hair cells expressing Myo7a. C-C`) High-magnification transverse showing Venus, Sox2, and Myo7a expression and the typical pseudostratified epithelial morphology of otic organoids. D-D` Low magnification projection view of the inside of a relatively large organoid stained with antibodies to Myo7a and the hair bundle protein Espin. E and F) Higher magnification views of the boxed regions indicated in D (inverted in E) showing the arrangement of Espin-labeled hair bundles on the apical surface of the epithelium.

### Venus expression in the developing and adult cochlea of *Fbxo2*^*VHC*^ mice

Two different *Fbxo2*^*VHC/WT*^ ES cell clones were used to generate two independent *Fbxo2*^*VHC/WT*^ knock-in mouse lines that showed identical expression patterns. All data reported here is derived from one of the two lines. Founder *Fbxo2*^*VHC/WT*^ mice were crossed to a ubiquitous zygotic Cre driver mouse line (Eiia-Cre, Lakso et al. (1996)) to remove the pgk-Neo cassette required for positive selection of ES cell clones (Fig. 1C). We examined Venus expression in developing and adult *Fbxo2*^*VHC/WT*^ mice and found that expression overlaps with Fbx2 immunoreactivity while also revealing more subtle expression level differences between cell types. Venus expression is moderate in the otic vesicle, where it overlaps with Fbx2 protein and the neurosensory domain marker Jag1 (Fig. 4A-A`). During cochlear morphogenesis, Venus expression is particularly intense throughout the floor of the developing cochlear duct, within the expression domain of Gata3 (Fig. 4B-B`), which is involved in specification of the prosensory domain (Luo et al., 2013). In the apex of the cochlea at E13.5, Venus expression spans the width of the cochlear floor and overlaps with the prosensory marker Sox2 (Fig. 4C-C`). A gradient was notable from the apex to the base where Venus expression was markedly lower on the medial side of the cochlea including the prosensory region and the greater epithelial ridge (Fig 4B-C`). Throughout embryonic and postnatal cochlear development this gradient of Venus and Fbx2 expression in the medial portion of the cochlea is maintained, so that the domain of Venus expression is wider in the apex and narrower and more restricted to the lateral cells in the base. This is markedly obvious in sections from E16.5 cochlea stained with antibodies to mark the organ of Corti and prosensory domain (Fig. 5A-E`). In the adult cochlea, Venus fluorescence is intense in organ of Corti hair cells and supporting cells, which also express Fbx2 protein (Fig 5.F-H`). Venus and Fbx2 are expressed strongly in all of the epithelial cell subtypes in the organ of Corti, including inner sulcus and outer sulcus supporting cell types as well as root cells located in the lateral extremity of the outer sulcus (Fig. 5F-G`). Lower levels of Venus and Fbx2 are present in spiral ligament fibrocytes of the lateral wall (Fig. 5F-G`). Within the sensory region of the adult organ of Corti, Venus expression is slightly lower in the outer hair cells compared to inner hair cells and supporting cells (Fig. 5H-H`).

**Figure 4.**
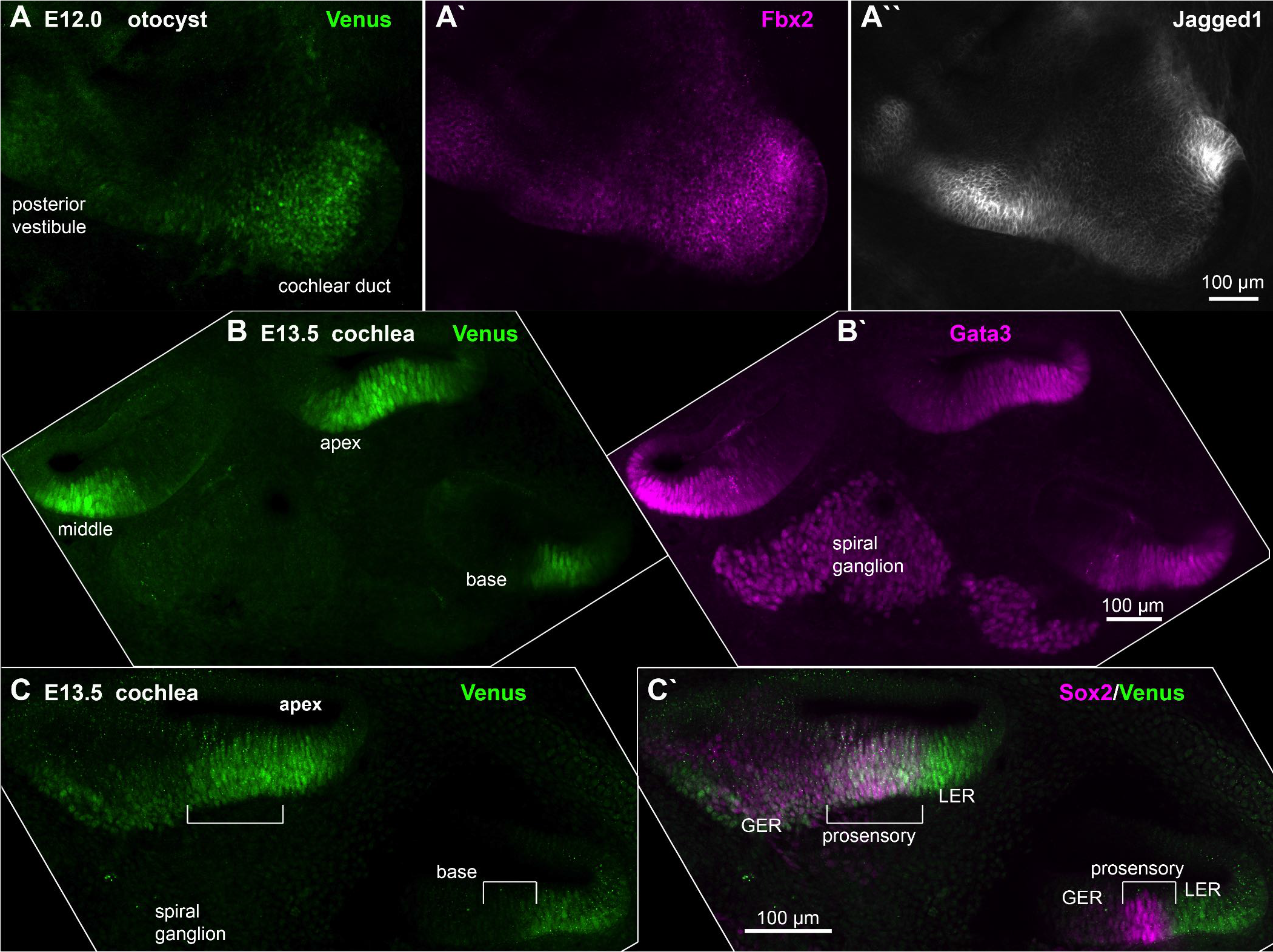
Venus expression in the developing otocyst of *Fbxo2*^*VHC/WT*^ heterozygous mice. Confocal micrographs of inner ear tissue processed for fluorescent immunolabeling with the indicated markers. A-A``) Individual imaging channel views of an E12.0 otocyst showing immunofluorescence signals for Venus (from the heterozygous VHC knock-in allele at the *Fbxo2* locus), Fbx2 (from the wild type *Fbxo2* locus) and Jag1, which marks prosensory/proneural domains. This sample was imaged from the medial surface of a whole-mount preparation and is shown as a projected z-stack. B-B`) Individual views of Venus and Gata3 immunofluorescence signals in an E13.5 cochlea vibratome section showing base, middle, and apical turns of the early cochlear duct. Venus expression in the floor of the cochlear duct is notably broad in the apex and narrow in the base, as shown by comparison to Gata3. C-C`) Similar section of E13.5 cochlea showing overlap of Venus and Sox2, which marks the prosensory domain at this stage. Abbreviations: GER, greater epithelial ridge; LER, lesser epithelial ridge

**Figure 5.**
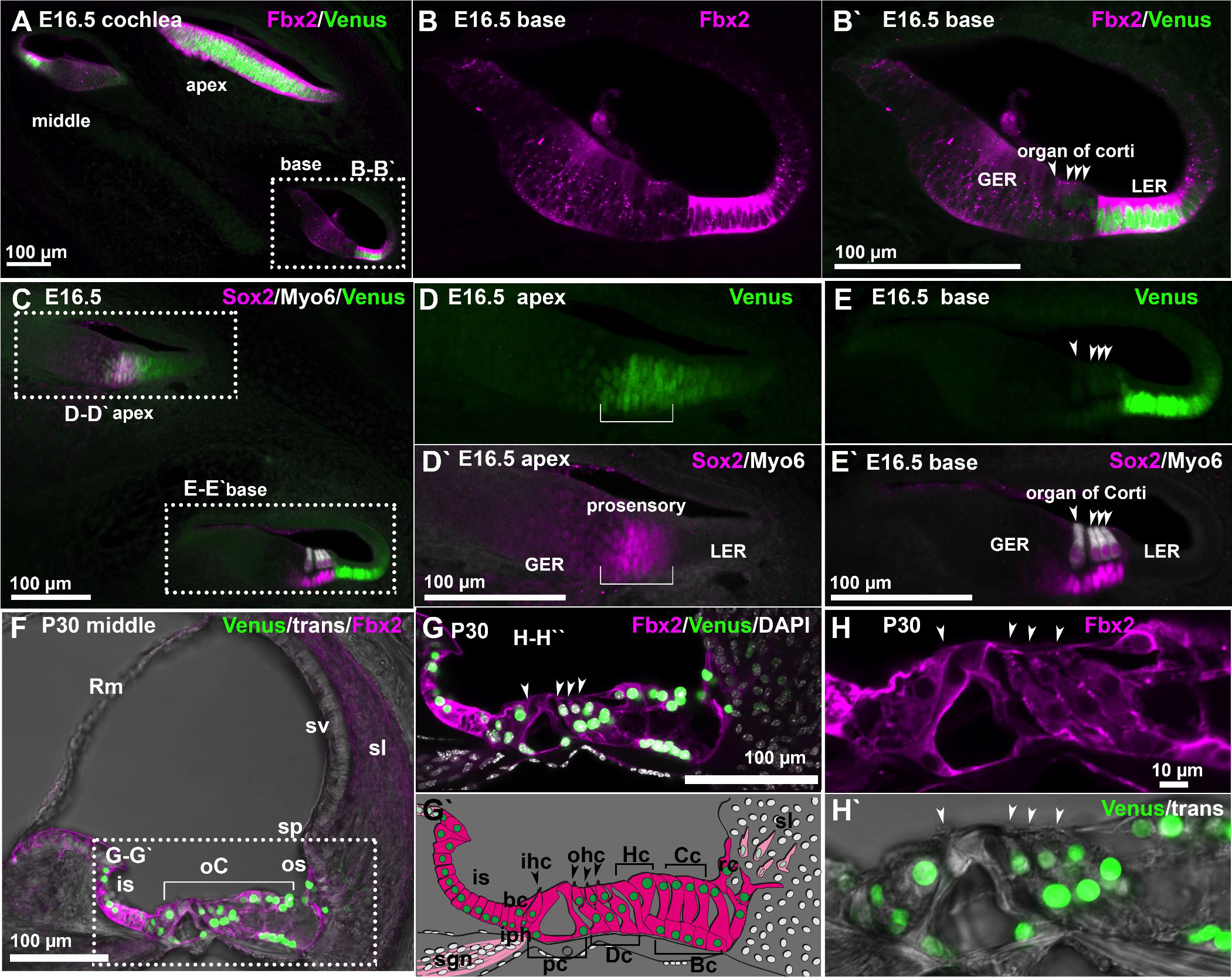
Venus Expression in the developing and adult organ of Corti of *Fbxo2*^*VHC/WT*^ heterozygous mice. Confocal micrographs of vibratome sections of cochlea from E16.5 (A-E`) and P30 (F-H`) *Fbxo2*^*VHC/WT*^ heterozygous mice processed for fluorescent immunolabeling with the indicated markers. A) In the E16.5 cochlea Fbx2 immunofluorescence overlaps with Venus in the floor of the cochlear duct, with more broad expression in the apex and a narrow domain in the base. B-B`) A view of the basal turn shows high expression of Fbx2 and Venus in the LER and moderate expression in the GER and organ of Corti. C-E`) Views of an E16.5 cochlea labeled with antibodies to the prosensory marker Sox2 and the hair cell marker Myo6. D-D`) Venus expression in the apex is found throughout the Sox2-expressing prosensory domain, prior to differentiation of hair cells in this region. E-E`) Myo6 signal is present in developing hair cells and Sox2 is expressed in developing supporting cells of the cochlea base at E16.5. Venus expression is high in the LER and low to moderate in the developing hair cells and Sox2-positive supporting cells. F) View of the scala media from the middle turn of the P30 cochlear duct shows high expression of both Fbx2 and Venus in the organ of Corti and the inner sulcus and outer sulcus. G) A higher magnification view of the boxed region in (F), shown with the nuclear marker DAPI illustrates the specific distribution of Fbx2 and Venus. G`) Schematized representation of Fbx2 and Venus expression patterns in the adult organ of Corti, with individual cell types indicated. H-H`) High magnification views of the organ of Corti show the distribution of Fbx2 in the cytoplasm of hair cells and supporting cells and expression of Venus in the nuclei of the same cells. Abbreviations: GER, greater epithelial ridge; LER, lesser epithelial ridge; Rm, Reissner’s membrane; sv, stria vascularis; sl, spiral ligament; sp, spiral prominence; is, inner sulcus; oC, organ of Corti; os, outer sulcus; bc, border cell; ihc, inner hair cell; ohc, outer hair cells; HC, Hensen’s cells; Cc, Claudius cells; rc, root cells; sgn, spiral ganglion nerve fibers; pc, pillar cells; Dc, Deiters cells; Bc, Boettcher cells.

### Tamoxifen-inducible Cre-recombination in the cochlea of *Fbxo2*^*VHC*^ mice

We tested Cre recombination in *Fbxo2*^*VHC*^ mice by crossing to the Ai14 Cre-reporter mouse (Madisen et al., 2010). Tamoxifen was administered as a single dose at various time points and samples were analyzed after 3 or more days for tdTomato-expressing cells. Cre recombination occurs sparsely in the cochlea when induced in mid-gestation embryos with a single dose of tamoxifen to the pregnant dam (Fig. 6A-C`). The frequency of tdTomato-expressing cells increases progressively at later stages of embryogenesis and with age in postnatal to adult animals (Fig. 6D-K). Tracing in the neonatal cochlea occurs sporadically in organ of Corti supporting cells and hair cells and more frequently in lateral epithelial ridge supporting cells (Fig 6E-H`). An increasing base to apex gradient of recombination frequency in the neonatal organ of Corti correlates with lower Venus expression levels in the base (Fig. 6 E-H`).

**Figure 6.**
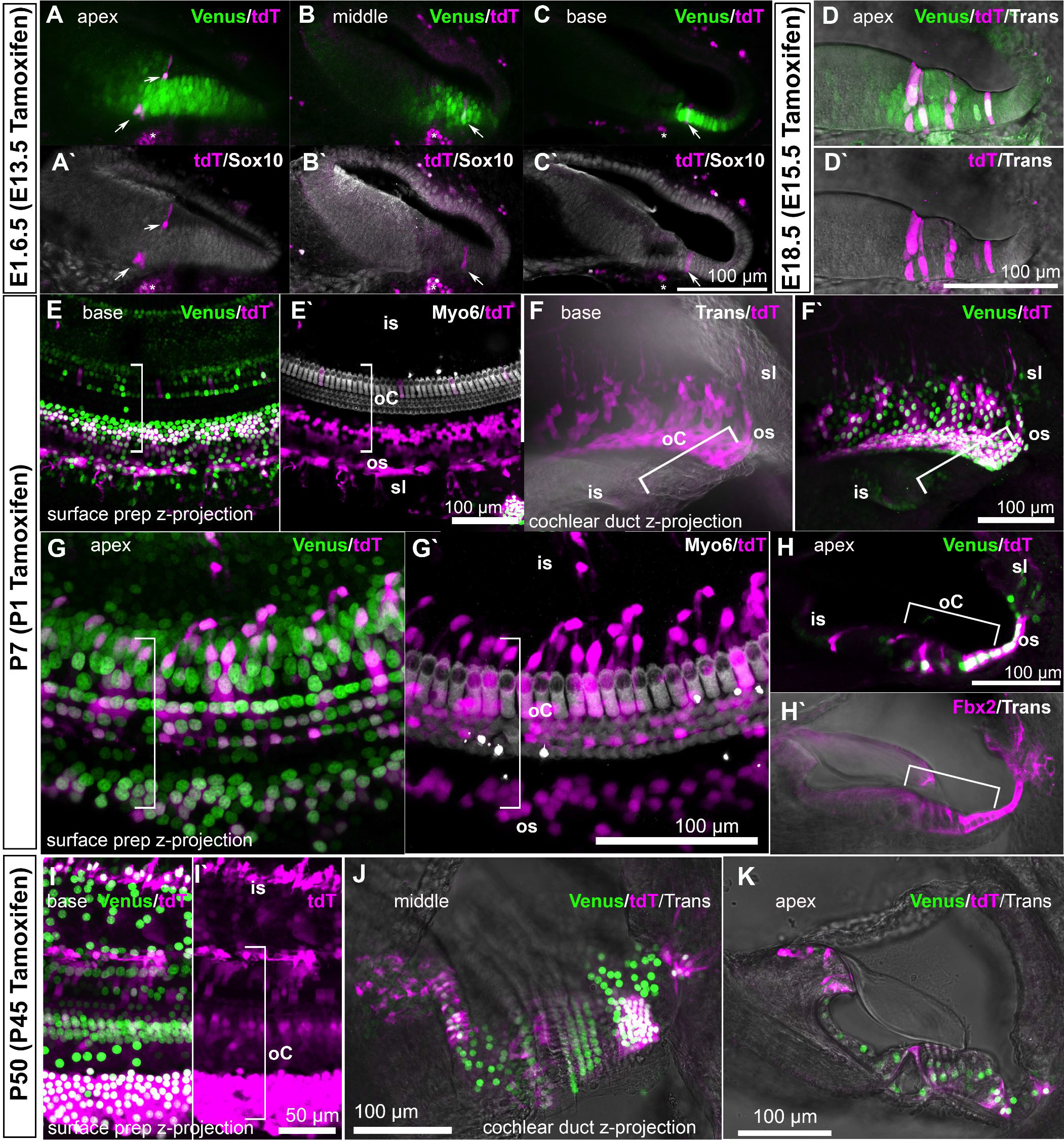
CreER recombination in the cochlea of *Fbxo2^VHC/WT^;Rosa26^Ai14/WT^* mice. *Fbxo2*^*VHC*^ mice were crossed to Ai14 reporter mice to generate compound heterozygotes. In these animals, Cre recombination is induced by Tamoxifen injection and cells that have undergone recombination express the fluorescent protein tdTomato (tdT). Tamoxifen was delivered via intraperitoneal injection to pregnant dams carrying embryos at E13.5 (A-C`) or E15.5 (D-D`), or to pups at P1 (G-H`) or adults at P45 (I-K).Animals were euthanized and collected at the indicated ages and cochleae were processed for either vibratome sections (A-D`, F-F`, H-H`, J, K) or whole-mount surface preparations (E-E`, G-G`, I-I`). Immunofluorescence labeling for Sox10 (cochlear duct epithelial cells, A-C`) or Myo6 (hair cells, E-E`, and G-G`) was performed on some samples as indicated. Venus and tdTomato and fluorescence were imaged directly in unstained samples (D-D`), (F-F`), (H-J). Abbreviations: is, inner sulcus; oC, organ of Corti; os, outer sulcus; sl, spiral ligament

### Cochlear degeneration in aged homozygous *Fbxo2*^*VHC/VHC*^ mice

The modified *Fbxo2*^*VHC*^ locus introduces a TGA stop codon after the VHC cassette and therefore is expected to be a null allele for Fbx2 (Fig. 1C). We found that homozygous *Fbxo2*^*VHC/VHC*^ mice exhibit age-related cochlear degeneration, which aligns with previous observations in Fbx2-null mice (Nelson et al., 2006). We compared cochleae of *Fbxo2*^VHC/WT^and *Fbxo2*^*VHC/VHC*^ littermates (FVB/NJ background) at eight months of age and performed immunohistochemistry for Myosin7a and Sox2 to label hair cells and adjacent supporting cells, respectively (Fig. S3). Cochleae of heterozygous animals in these aged animals were indistinguishable from P30 samples and did not display any loss of cells or abnormal cell morphologies (Fig. S3A-B`, compare to Fig. 5F). Homozygous *Fbxo2*^*VHC/VHC*^ cochleae were found to have abnormal morphology consistent with degeneration of hair cells and supporting cells. This phenotype was present throughout the length of the cochlea, but was more severe in the base. The middle turns displayed morphological abnormalities of hair cells and supporting cells while the base of the cochlea appeared as a flattened epithelium devoid of hair cells (Fig. S3C-F`). Even in areas of severe degeneration, some Venus-expressing cells remain and appear to have intact nuclei (Fig. S3F).

### *Fbxo2*^*VHC*^ reporter activity marks a subset of vestibular hair cells

In the utricle, Venus is initially detected at low levels in some Sox2-labeled progenitors/supporting cells (E13.5, Fig. S4A). At E16.5, some hair cells express Venus at higher levels, while expression in the supporting cell layer remains low (Fig. S4B-D). In postnatal and adult vestibular organs, Venus is bright in nuclei of cells located almost exclusively in the peripheral/extrastriolar zones (Fig. 7). In the mammalian macular organs, the utricle and saccule, the striolar region of sensory epithelium is immediately adjacent to the line of hair cell polarity reversal and displays unique features including oncomodulin (Ocm)-positive hair cells and afferent dendrites exhibiting primarily calyx-only morphology and calretinin expression (Desai et al., 2005b; Hoffman et al., 2018; Simmons et al., 2010). The saddle-shaped semi-circular canal cristae exhibit a central zone with features very similar to the macular striola (Desai et al., 2005a). Close examination and immunohistochemical analysis of *Fbxo2*^*VHC/WT*^ macular organs show that the Venus-expressing cells are Myosin6-positive hair cells located outside of the striolar zone (Fig. 7A-B``). Fbx2 immunoreactivity is present in Venus-positive cells and appears to be enriched in perinuclear cytoplasm (Fig. 7C).

**Figure 7.**
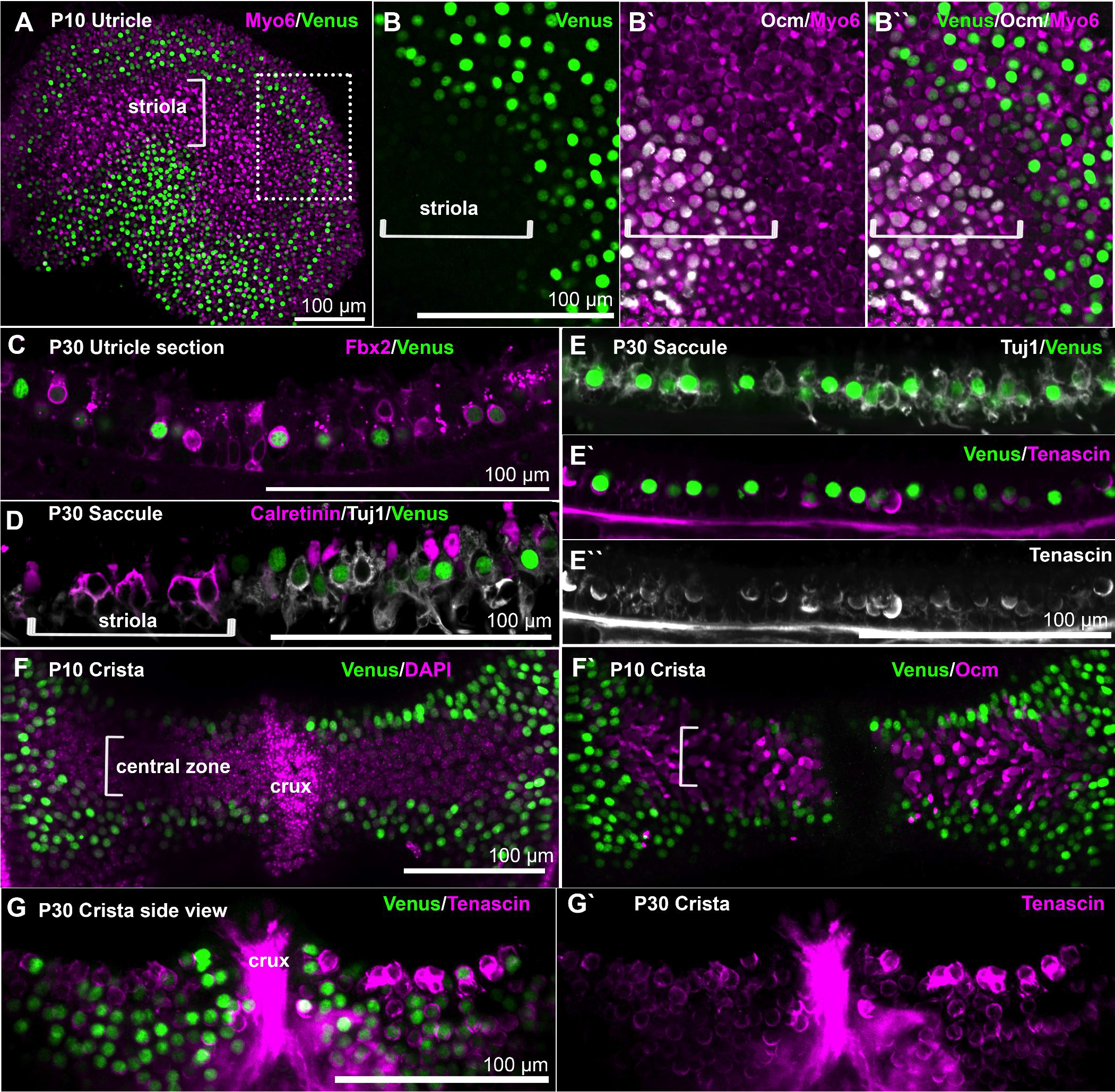
*Fbxo2*^*VHC*^ expression marks extrastriolar/non-central type I vestibular hair cells. Confocal micrographs of P10 (A-B``, F-F`) and P30 (C-E``, G-G`) vestibular organs from heterozygous *Fbxo2*^*VHC*^ mice processed for fluorescent immunolabeling with the indicated markers, as whole-mount (A-B``, F-G`) or vibratome sections (C-E``). (A-B`) Immunofluorescence for Venus, Myo6 and Ocm in a P10 utricle shows Venus expression in nuclei of Myo6-positive hair cells in extrastriolar regions, but not in Ocm-positive hair cells of the striolar zone. C) Venus expression correlates with Fbx2 immunofluorescence in a P30 utricle vibratome section. D) Vibratome section of the P30 saccule showing calretinin expression in compound calyx nerve endings of the striolar zone (bracket) as well as type II hair cells in the extrastriolar zone (right); signals for Venus and Tuj1 are overlaid as indicated. Venus-positive nuclei are seen in cells associated with calyx innervation typical of type I hair cells, but only those in extrastriolar regions. E-E`) Image of a vibratome section from an extrastriolar region of P30 saccule showing Venus expression in nuclei of the hair cell layer, surrounded by simple calyx nerve endings (Tuj1) and crescent-shaped labeling of Tenascin, an extracellular matrix protein specifically found in calyxes of vestibular sensory epithelia. F-F`) Z-projection top-down views of a whole-mount P10 crista show the distribution of Venus-positive nuclei in the non-central zone of the sensory epithelia, overlaid with the nuclear stain DAPI (F) or immunofluorescence signal for Ocm (F`). G-G`) Z-projection side views of a P30 crista showing Venus expression in cells associated with Tenascin-labeled calyxes, specific to type I hair cells.

Mammalian vestibular organs have two major types of hair cells, types I and II, present in similar numbers (for review see Eatock and Songer (2011)). Type I vestibular hair cells are characterized by calyx afferent innervation while type II hair cells are innervated by bouton nerve endings (Goldberg et al., 1992; Lysakowski and Goldberg, 1997). In mature vestibular organs, calretinin is expressed in type II hair cells but not type I hair cells (Dechesne et al., 1994; Li et al., 2008). Vestibular sensory epithelia costained with calretinin and the neuronal marker Tuj1 showed that Venus-positive cells are calretinin negative and are associated with calretinin-negative calyxes in the extrastriolar region, suggesting that Venus expression is specific to extrastriolar type I hair cells (Fig. 7D). To further validate this finding, we immunolabeled for tenascin, an extracellular protein specifically found in the region between type I vestibular hair cells and their surrounding afferent nerve calyxes (Lysakowski et al., 2011; Warchol and Speck, 2007). We found that each Venus-positive cell is surrounded by a crescent of tenascin immunoreactivity (Fig. 7E-E``). The *Fbxo2*^*VHC*^ reporter shows a similar pattern of expression in semicircular canal cristae to that observed in macular organs. Venus is expressed in hair cell nuclei in the peripheral zones but is absent from the central zone (Fig. 7F). As in the utricle, Venus expression is juxtaposed with the striolar hair cell marker oncomodulin (Fig. 7F`). Moreover, Venus-labeled nuclei are nearly always associated with rings or crescents of tenascin immunoreactivity in the crista (Fig. 7G-G`). We conclude that *Fbxo2*^*VHC*^ specifically labels populations of type I hair cells found in the extrastriolar zones of the maculae and the peripheral zones of the cristae.

Cre recombination in *Fbxo2*^*VHC*^ vestibular organs is robust and consistent with the Venus expression observed in developing and adult organs (Fig. S5). When tamoxifen was delivered to pregnant dams at E13.5 no Cre recombination was observed in the vestibular organs (data not shown). An E15.5 dose of tamoxifen resulted in occasional recombination in hair cells in the peripheral/extrastriolar zones when samples were collected at E18.5 (Fig. S5A-A`). Moreover, labeled cells, which were presumably Venus-positive at the time of tamoxifen delivery, continue to express Venus at the time of analysis (Fig. S5A-A`). Recombination occurs in large numbers of extrastriolar/non-central hair cells following injection of tamoxifen to newborn mice (Fig. S5B-F``). At six days following tamoxifen injection most of the Venus-expressing cells in the utricle are labeled with tdTomato and vice versa (Fig. S5C-C``). Fbx2 protein immunoreactivity is visible and appears enriched in the cells labeled with Venus and tdTomato in both macular and ampullar organs (Fig. S5E-E` and F-F``).

We examined vestibular organs of homozygous *Fbxo2*^*VHC/VHC*^ mice at eight months of age to determine if there are any signs of degeneration in the vestibule, given the progressive cochlear degeneration phenotype observed in aged Fbx2-null mice. No indication of gross morphological abnormalities or cell loss was observed in vestibular organs of homozygous animals (Fig. S6). Distribution of Venus-positive cells and calyxeal nerve endings also appeared normal in the utricle and crista. Neither did we observe any circling or whirling behavior in *Fbxo2*^*VHC/VHC*^ animals, consistent with previously reported analysis of Fbx2-null animals (Nelson et al., 2006).

### Cerebellar granule cells express the *Fbxo2*^*VHC*^ reporter

We examined organism-wide Venus expression and Cre reporter activity in whole embryos and vibratome sections from E10.5 to birth to identify other sites of substantial *Fbxo2*^*VHC*^ expression besides the inner ear. This analysis identified cerebellar granule cells and their precursors as the only other population exhibiting expression of the reporter at levels comparable to the inner ear. In sagittal vibratome sections of the head at E16.5, bright Venus expression is present in the external granular layer of the cerebellum (Fig. S7A-C). The external granular layer, or external germinal zone, is a temporary population of proliferating progenitors located at the subpial surface of the cerebellum (for review see (Marzban et al., 2014)). This population is Atoh1-dependent and is derived from the rhombic lip at the dorsal edge of the fourth ventricle between E12.5 and E17 (Machold and Fishell, 2005; Wang and Zoghbi, 2001). In lineage tracing experiments, injection of tamoxifen at E13.5 resulted in moderate Cre recombination within this population, and no other sites of the brain (Fig. S8A-A``).

The vast majority of cells from the external granular layer produce cerebellar granule cells during the postnatal period in rodents. During rodent postnatal development, granule cells differentiate and migrate rostrally across the Purkinkje cell layer to form the granular layer, the innermost layer of the mature cerebellum. Tamoxifen administration at P1 resulted in labeled cells in the external granular layer and in cells that appear to be migrating to the inner granular layer (Fig. S8B-B``). In adults, Venus fluorescence is bright in the granular layer in all lobes of the cerebellum (Fig. S7D-D`). There are also some other regions of the adult brain that harbor Venus expressing cells, including the pons, medulla and hippocampus, but the intensity of fluorescence is lower and expression is sparse in these areas (Fig. S7D-D`). Closer examination of the cerebellum revealed bright Venus expression in densely packed nuclei of the granular layer, which is bordered by the Purkinje layer, labeled with calbindin (Fig. S7E-E`). Venus-expressing cells are positive for calretinin (Fig. S7F-F`), which labels cerebellar granule cells in adult rodents (Resibois and Rogers, 1992). Cerebellar tissue from eight-month old *Fbxo2*^*VHC/WT*^ and littermate *Fbxo2*^*VHC/VHC*^ mice were stained with antibodies to calbindin and calretinin to visualize the Purkinje cells and granule cells, respectively (Fig. S7G-H), and no abnormalities were observed.

## DISCUSSION

Genetically engineered mouse models including gene targeted and transgenic reporters are essential tools of discovery in mammalian developmental biology (Bouabe and Okkenhaug, 2013). Specificity and utility of a transgenic model depends on two main factors. First, a tissue or cell-specific promoter/ locus is needed to express genes either consistently or transiently in the organ or cell lineage of interest. The second factor is the transgene product that is expressed under control of the tissue or cell-specific promoter; this might be a reporter protein or a more complex construct. The number of various promoter and reporter combinations is ultimately dependent on the research area, but usually restrictive so that it is often necessary to cleverly combine existing mouse models to address specific biological questions. Here, we introduce a combination of a highly specific otic lineage locus with a multicistronic reporter/effector cassette that enables a wide variety of functional experiments. We provide a toolbox of different reporter multicistronic cassettes that all have been tested, which are freely available to be combined with any gene provided by the user.

Moreover, we have thoroughly characterized the otic lineage-specific *Fbxo2*^*VHC/WT*^ ES cell line and derived transgenic mouse lines to showcase the various uses of our toolbox. The combination of three distinct functional proteins confers a variety of useful features. The H2B fusion to a bright fluorescent protein is not only a readout of gene expression, but can be used for live imaging of cell behavior and chromatin dynamics, as well as a means to sort live cells or nuclei via flow cytometry. HygroR confers drug resistance that may be used to select for reporter-expressing cells in culture, which can be applied to cell line development, directed differentiation or lineage reprogramming. Expression of CreER enables drug-inducible recombination for conditional deletion of LoxP-flanked genes or conditional activation of Cre-inducible transgenes, such as for lineage tracing or gain-of-function studies. With respect to *Fbxo2*^*VHC/WT*^ reporter expression, we found high correlation between *Fbxo2* mRNA, Fbx2 protein, and VHC reporter expression during development, in neonatal and adult mice, as well as in ES cell-derived otic organoids. Figure 8A summarizes our findings for *Fbxo2*^*VHC*^ reporter activity in mice as indicated by Venus fluorescence and Cre recombination. Cell populations with high Venus expression showed higher rates of Cre recombination as well as higher levels of Fbx2 immunoreactivity. Fbx2 and the *Fbxo2*^*VHC*^ reporter both show a steady increase in expression levels during later development and maturation, so the highest levels of expression are found in mature cells. The locations of highest expression in the adult include the organ of Corti and the granular layer of the cerebellum. In the late embryonic and neonatal cochlea, the *Fbxo2*^*VHC*^ reporter revealed a gradient with highest expression in the lateral supporting cells. In vestibular organs, expression of the *Fbxo2*^*VHC*^ reporter is highest in a unique subset of type I hair cells, some of which appear to be specified during embryonic development as indicated by lineage tracing and retained Venus expression. Overall expression patterns of the *Fbxo2*^*VHC*^ reporter are consistent with Fbx2 protein expression levels reported here as well as in previous work (Atkin et al., 2014; Hartman et al., 2015; Nelson et al., 2007). As well, observed *Fbxo2*^*VHC*^ reporter expression aligned well with *Fbxo2* transcriptional expression profiles from existing RNA-Seq studies (Fig. 8B). Analysis of the transcriptional distribution of *Fbxo2* in cellular subpopulations of the cochlea and utricle of P1 mice (data from Burns et al. (2015)) shows that cochlear lateral supporting cells exhibit significantly higher expression than other cochlear cell populations and reveal a subpopulation of utricle hair cells having elevated *Fbxo2* levels, consistent with the *Fbxo2*^*VHC*^ reporter. Notably, a cross-species comparison of *Fbxo2* expression in adult vertebrate tissues indicates a highly conserved enrichment of *Fbxo2* mRNA in brain and cerebellum.

**Figure 8.**
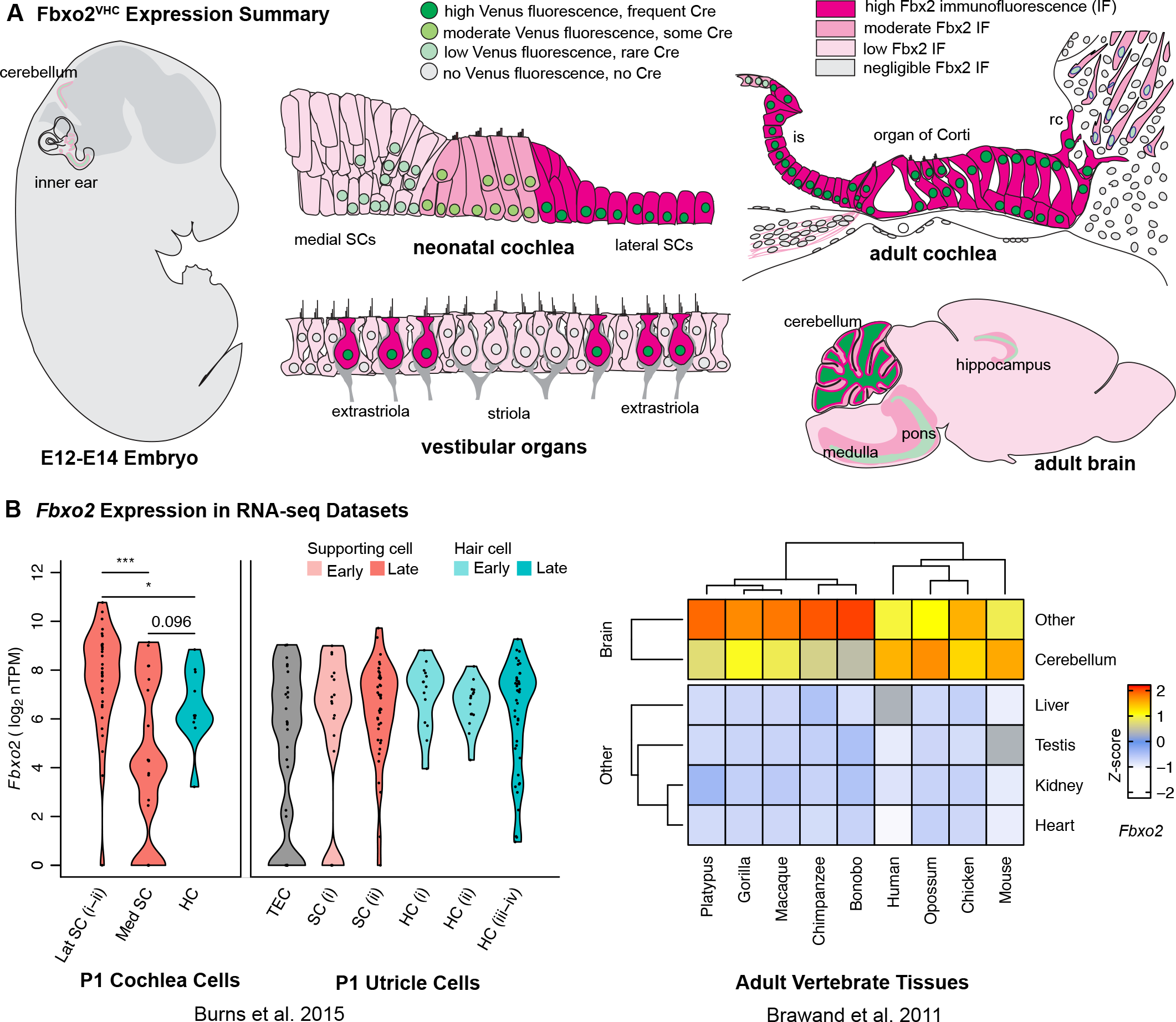
*Fbxo2*^*VHC*^ and Fbx2 Expression Patterns are Consistent with *Fbxo2* Expression Trends in RNA-seq Datasets. A) Schematic of *Fbxo2*^*VHC*^ reporter activity and Fbx2 protein levels in developing and adult mouse tissues. B) Expression of *Fbxo2* in single cells from P1 mouse cochlea (left panel) and utricle (right panel) tissue. Cells were grouped per tissue according to Burns et al. (2015). Violin plots show vertically mirrored *Fbxo2* expression density curves per group; individual data points denote *Fbxo2* expression in single cells. Abbreviations: TEC = transitional epithelial cells, SC = supporting cells, HC = hair cells, Lat = lateral, Med = medial; nTPM = normalized transcripts per million; * combined Likelihood Ratio Test p < 0.05, *** p < 0.001. B) Heat map of *Fbxo2* expression in different tissues and organisms as measured by Brawand et al. (2011); expression is scaled per organism to show differential *Fbxo2* expression per species (Z-score).

In the organoid system we show that 1) the *Fbxo2*^*VHC*^ reporter is effective for delineating the otic lineage, 2) Venus expression is found in both hair cells and supporting cells at moderate to high levels, and 3) elevated Venus expression can be found in some patches of hair cells and supporting cells. Interestingly, this pattern is distinct from that of the native vestibular organs in *Fbxo2*^*VHC*^ mice, in which expression in supporting cells is quite low and only a subset of hair cells express moderate to high Venus. This was somewhat unexpected given that inner ear organoids produced from other ES cell lines with the same protocol have been shown to contain hair cells that are morphologically and functionally similar to native vestibular hair cells (Liu et al., 2016). In *Fbxo2*^*VHC*^ organ of Corti, expression of Venus is found in all epithelial cells, and is elevated in the supporting cells of the lateral domain during embryonic and postnatal development. Expression of Venus in what appears to be all supporting cells and hair cells in organoids, indicates that regulation of *Fbxo2* in these ES cell-derived sensory epithelia may not be entirely recapitulating the signaling of vestibular organogenesis. The presence of patches of higher expressing cells within organoids suggests that *Fbxo2* may be affected by signaling domains that may occur in the aggregate cultures. Signaling from BMP, Wnt, and Notch regulate mediolateral patterning of the cochlea and likely affect the gradient of Fbx2/*Fbxo2*^*VHC*^ in the organ of Corti (Munnamalai and Fekete, 2016; Ohyama et al., 2010). Further investigation into the properties of otic organoid cells and signaling can be facilitated by use of the *Fbxo2*^*VHC*^ reporter. As well, *Fbxo2*^*VHC*^ ES cells may be applied to ES cell models of cerebellar development (Salero and Hatten, 2007; Srivastava et al., 2013).

In conclusion, the *Fbxo2*^*VHC*^ reporter system is a versatile tool for the study of developmental biology of the inner ear and the cerebellum. These validated mouse and ES cell models complement each other and can be used in parallel to take advantage of the relative strengths of each system. The multicistronic reporter approach and spectrally distinct reporter toolbox developed here is now available as a resource for the developmental biology community.

## MATERIALS AND METHODS

### Cloning

Multicistronic cassettes were constructed using enzymatic assembly of overlapping DNA fragments (Gibson assembly, (Gibson, 2011)). Source plasmids for cloning are listed in supplemental methods. To generate the Fbxo2-VHCpgkNeoDTA targeting construct (Figure 1 C), homology arms flanking Exon 1 of *Fbxo2* were amplified from E14tg2a ES cell genomic DNA using high fidelity PCR. Further details are provided in supplemental methods.

### Generation of *Fbxo2*^*VHC*^ mouse ES cells and mice

ES cell manipulation and selection was performed by the Stanford Transgenic, Knockout and Tumor Model Center (TKTMC). Further details are provided in supplemental methods.

### *Fbxo2*^*VHC*^ mouse ES cell culture and generation of otic organoids

*Fbxo2*^*VHC/WT*^ ES cells were expanded from the same clone used to produce the mouse line described in this report. ES cells were cultured without feeders in “2iLif” conditions (leukemia inhibitory factor (LIF) in serum-free N2B27 supplemented with MEK inhibitor PD0325901 (1 μM) and GSK3 inhibitor CHIR99021 (3 μM) (Ying et al., 2008)). Inner ear sensory epithelia were differentiated from *Fbxo2^VHC/WT^* ES cells in serum-free 3D aggregate culture as previously described with only very minor modifications (Koehler and Hashino, 2014; Koehler et al., 2013). Briefly, *Fbxo2*^*VHC/WT*^ ES cells in 2iLif were washed well in DMEM/F12 then dissociated with Accutase (Stem Cell Technologies) and diluted to a concentration of 30,000 cells per ml in ectodermal differentiation medium and aliquoted at 100μl per well in 96-well ultra low attachment plates to form aggregates. Matrigel was added 24 hours later on day 1, BMP-4 and SB-431542 were added on day 3, and FGF-2 and LDN-193189 (FGF/LDN) were added on day 4 to 5. On day 8, aggregates were transferred to long-term culture to be maintained until endpoint analysis. As described by Koehler and Hashino (2014), the timing of FGF/LDN was an important factor in optimizing the protocol; application on day 4.0 to 4.25 produced higher rates of organoids than did application on 4.5 to 5.0 in *Fbxo2*^*VHC/WT*^ ES cells.

### Fluorescent immunohistochemistry and imaging

Organoids, whole embryos, or dissected tissues were processed either whole or in vibratome sections. Samples were permeabilized with 2% Triton X-100 in PBS for one hour at room temperature then incubated in blocking solution (10% fetal bovine serum, 0.5% Triton X-100, in PBS) overnight at 4°C. Primary antibodies (see Supplemental Methods for list of antibodies) were diluted in blocking solution and samples were incubated for two to three nights at 4°C with rocking. Samples were washed then incubated in species-specific Alexafluor 488-/568-/ or 647-conjugated secondary antibodies (Life Technologies) diluted in blocking solution. Vibratome sections were incubated for at least one hour in 50% glycerol in PBS then mounted in the same on a glass concavity slide and coverslipped for imaging. Aggregates and whole mount tissue specimens were optically cleared by incubating at 4°C with rocking in Scale/A2 (Hama et al., 2011) for at least 5 days, with three changes of the Scale/A2 solution. Cleared samples were mounted in Scale/A2 between glass coverslips in imaging chambers consisting of 2-3 stacked 0.5 mm adhesive silicone spacers, modified from SecureSeal Hybridization
Chambers (Grace Biolabs). Confocal fluorescent imaging was performed with upright Zeiss LSM700 or LSM880 confocal microscopes running Zen software and using 10X, 20X, or 40X objectives. ImageJ/ FIJI was used for image analysis and export of z-stack data to video (AVI). Adobe CS6 (Adobe Systems) was used for processing images into figures. Keynote (Apple) was used to combine video files and generate movies with titles and annotations. Further details are provided in Supplemental Methods.

## ACKNOWLEDGMENTS

We thank Andy Groves for consulting on this project, Hong Zeng for gene targeting support, David Bredt for Fbx2 antibodies, Robert Campbell and Michael Davidson for fluorescent protein reagents and advice, and members of the Heller lab for constructive comments on the manuscript. Imaging was performed at the Stanford OHNS Imaging Core Facility and supported by core grant P30DC010363. ES cell manipulation and blastocyst microinjection were performed by the Stanford University Transgenic, Knockout and Tumor Model Center.

## DATA AVAILABILITY/ RESOURCE SHARING

Plasmid DNA sequences and constructs are available from the nonprofit plasmid repository Addgene (www.addgene.com, Cambridge MA). Fbxo2^VHC^ mice and ES cells are available from the Mouse Mutant Resource & Research Centers supported by NIH (MMRRC, www.mmrrc.org).

## AUTHOR CONTRIBUTIONS

B.H.H. and S.H. conceived the study, interpreted the results, and wrote the manuscript. B.H.H. designed and generated the DNA constructs and performed the experiments. R. B. and S.K. performed some of the ES cell differentiation experiments, immunolabeling, and imaging. D. C. E. performed analysis of transcriptional expression profiles and statistics. M.S. assisted with cloning and cell culture. All authors provided feedback and approved the final manuscript.

